# Glycolytic inhibitor 2-deoxyglucose prevents cortical hyperexcitability after traumatic brain injury

**DOI:** 10.1101/479782

**Authors:** Jenny B. Koenig, David Cantu, Cho Low, Farzad Noubary, Danielle Croker, Michael Whalen, Dong Kong, Chris G. Dulla

**Affiliations:** Department of Neuroscience, Tufts University School of Medicine, Boston, MA, USA.; Neuroscience Program, Tufts University Sackler School of Biomedical Sciences, Boston, MA, USA.; Cellular, Molecular, and Developmental Biology Program, Tufts University Sackler School of Graduate Biomedical Sciences, Boston, MA, USA.; Department of Health Sciences, Bouvé College of Health Sciences, Northeastern University, Boston, MA, USA.; Neuroscience Center, Harvard Medical School, Massachusetts General Hospital, Charlestown, MA, USA.; Department of Pediatrics, Harvard Medical School, Massachusetts General Hospital, Boston, MA, USA.

## Abstract

Traumatic brain injury (TBI) causes cortical dysfunction and can lead to post-traumatic epilepsy. Multiple studies demonstrate that GABAergic inhibitory network function is compromised following TBI, which may contribute to hyperexcitability and motor, behavioral, and cognitive deficits. Preserving the function of GABAergic interneurons, therefore, is a rational therapeutic strategy to preserve cortical function after TBI and prevent long-term clinical complications. Here, we explored an approach based on the ketogenic diet, a neuroprotective and anticonvulsant dietary therapy which results in reduced glycolysis and increased ketosis. Utilizing a pharmacologic inhibitor of glycolysis (2-deoxyglucose, or 2-DG), we found that acute *in vitro* glycolytic inhibition decreased the excitability of excitatory neurons, but not inhibitory interneurons, in cortical slices from naïve mice. Employing the controlled cortical impact (CCI) model of TBI in mice, we found that *in vitro* 2-DG treatment rapidly attenuated epileptiform activity seen in acute cortical slices 3-5 weeks after TBI. One week of *in vivo* 2-DG treatment immediately after TBI prevented the development of epileptiform activity, restored excitatory and inhibitory synaptic activity, and attenuated loss of parvalbumin-positive inhibitory interneurons. In summary, inhibition of glycolysis with 2-DG may have therapeutic potential to restore network function following TBI.

**One Sentence Summary:** Following traumatic brain injury in mice, *in vivo* treatment with the glycolytic inhibitor 2-deoxyglucose prevented cortical network pathology including cortical hyperexcitability, changes in synaptic activity, and loss of parvalbumin-expressing GABAergic interneurons.

## Introduction

Traumatic brain injury (TBI) is a leading cause of death and disability worldwide and can lead to motor, behavioral, and cognitive losses (*1*) as well as post-traumatic epilepsy (PTE) (*2–4*). Patients who develop PTE show higher mortality than TBI patients without epilepsy, and many PTE patients are refractory to the currently available anticonvulsive therapies (*2, 4*). Multiple molecular- and cellular-level changes that occur after TBI may play an important role in the pathophysiology of network dysfunction, PTE, and other TBI-induced pathologies. Aberrant electrical activity (*5, 6*), indiscriminate release of neurotransmitters (*7*), and altered cerebral glucose utilization (*8, 9*) all occur after TBI. Additionally, compromised cerebral blood flow, disruption of the blood-brain-barrier (*10, 11*), increased oxidative stress, and mitochondrial dysfunction (*12*) create a complex set of neurometabolic disruptions after injury (*1, 2, 13*). Together, these changes can disturb synaptic function, cause hard-wired remodeling of neuronal connectivity, and lead to circuit dysfunction and PTE.

Many studies suggest that a key step in developing network dysfunction after injury is the loss of inhibitory GABAergic interneurons, a diverse group of neurons that powerfully constrain cortical excitation. Disruption of inhibitory network activity can lead to motor dysfunction, cognitive losses, and epilepsy (*14–18*). The most abundant subtype of cortical interneurons express the calcium-binding protein parvalbumin (PV), fire action potentials at a high frequency (fast-spiking), and provide perisomatic inhibition of excitatory cortical pyramidal neurons (*19*). PV+ interneuron loss, atrophy, or dysfunction has been shown in the controlled cortical impact (CCI) (*20*), lateral fluid percussion (*14*), partially isolated neocortex (“undercut”) (*21, 22*), and weight drop (*23*) models of TBI. Importantly, human post-mortem studies also show that PV-expressing interneurons are specifically lost following TBI (*24*). The mechanisms by which TBI leads to interneuron loss are largely unknown. Therefore, it has been difficult to develop therapeutic interventions capable of preserving inhibitory network function following TBI.

We believe that the aberrant neuronal activity that occurs acutely after TBI is a key contributor to long-term cortical network dysfunction. In support of this idea, early seizure activity (and other EEG abnormalities) are associated with poor clinical outcomes and higher risk of developing post-traumatic epilepsy (*5, 6, 25*). Preclinical data from animal models of epilepsy and brain injury demonstrate that early uncontrolled excitation contributes to later loss of GABAergic interneurons and development of inhibitory circuit dysfunction (*26–29*). Following TBI in both humans and animal models, this aberrant electrical activity occurs during periods of abnormally elevated glycolytic activity (*30–35*). We hypothesize that reducing hyper-glycolysis will attenuate injury-induced increases in excitatory synaptic activity, reduce losses of PV+ interneurons, and ultimately prevent inhibitory network dysfunction.

2-deoxyglucose (2-DG) is a glucose analog that competitively inhibits hexokinase (the rate-limiting enzyme in glycolysis) (*36–38*). Multiple studies demonstrate that glycolytic inhibition can be anticonvulsive in both *in vitro* and *in vivo* models of epilepsy (*39, 40*). 2-DG pre-treatment also prevents kainic acid-induced hippocampal cell loss in rats (*41*) and can reduce seizure severity (*42, 43*). While 2-DG has been previously explored as an anticonvulsant therapy, it has not yet been utilized as a disease-modifying therapy to prevent epileptogenesis after TBI.

In this study, we tested whether 2-DG can reduce TBI-induced hyperexcitation and attenuate the development of inhibitory network dysfunction after TBI. We utilized the controlled cortical impact (CCI) model of focal brain contusion, which results in significant cell loss, network hyperexcitability, and spontaneous behavioral and electrographic seizures (*44*). Our study shows that acute 2-DG treatment of cortical brain sections attenuated hyperexcitability in the injured brain. In addition, glycolytic inhibition reduced excitatory, but not inhibitory, neuronal excitability. Finally, *in vivo* treatment with 2-DG for one week following CCI prevented the development of epileptiform activity and reduced the loss of parvalbumin-expressing interneurons. Together, these studies support the potential application of glycolytic inhibitors to reduce cortical hyperexcitability after TBI.

## Results

### In vitro 2-DG treatment decreases the excitability of excitatory neurons, but not inhibitory interneurons

2-DG has been previously shown to attenuate epileptiform activity in multiple *in vitro* and animal models of epilepsy (*39, 40, 45*). Pharmacological inhibition of glycolysis can directly suppress synaptic transmission, reduce membrane resistance (*46*), and broaden the action potential (AP) waveform (*47*). It is unknown, however, if glycolytic inhibition has differential effects on the excitability of glutamatergic excitatory neurons and GABAergic inhibitory interneurons. Therefore, we assessed the effects of 2-DG on both cell types in naïve (uninjured) cortex. Whole-cell recordings were established in acute cortical brain slices from adult (>P28) animals of both sexes. Recordings were from layer V pyramidal neurons or fast-spiking PV+ GABAergic interneurons (identified by GFP expression in *G42* mice (*48*)). In current clamp mode, hyperpolarizing and depolarizing steps were injected in the presence of synaptic receptor blockers (10 µm CPP, 20 µm DNQX, and 10 µm GABAzine) and the number of evoked action potentials (APs) was quantified. The results were compared between baseline aCSF (containing 10 mM glucose) and following 10 minutes of slice perfusion with 2-DG-aCSF (8 mM 2-DG, 2 mM glucose). 2-DG-aCSF maintained the same solution osmolarity as baseline aCSF (10 mM total saccharide), while limiting the glucose availability to concentrations similar to those present in mouse or human cerebrospinal fluid (2 mM, (*49*)). In the presence of 2-DG, excitatory layer V pyramidal neurons fired significantly fewer APs at a given level of current injection (Figure 1A, 1C; Linear Mixed Model (LMM) – see Table S1 for all LMM results, fixed effect: interaction of current injection and 2-DG treatment; N = 14 cells from 9 animals; t = −4.129) while GABAergic interneurons were not affected (Figure 1B, 1D; LMM, not significant; N = 8 cells from 7 animals; t = 0.999). Consistent with reduced excitability of glutamatergic excitatory neurons following 2-DG perfusion, rheobase (current injection required to elicit the first AP) was increased in excitatory neurons but was not affected in GABAergic interneurons (Figure 1E; LMM, fixed effect: cell type; t = 2.184). The decrease in the intrinsic excitability of excitatory neurons was associated with a decrease in membrane resistance which was not observed in GABAergic interneurons (Figure 1F; LMM, fixed effect: cell type; t = −2.703). No change in resting membrane potential was observed in either cell type upon application of 2-DG (Figure 1G; LMM, not significant).

**Figure 1:**
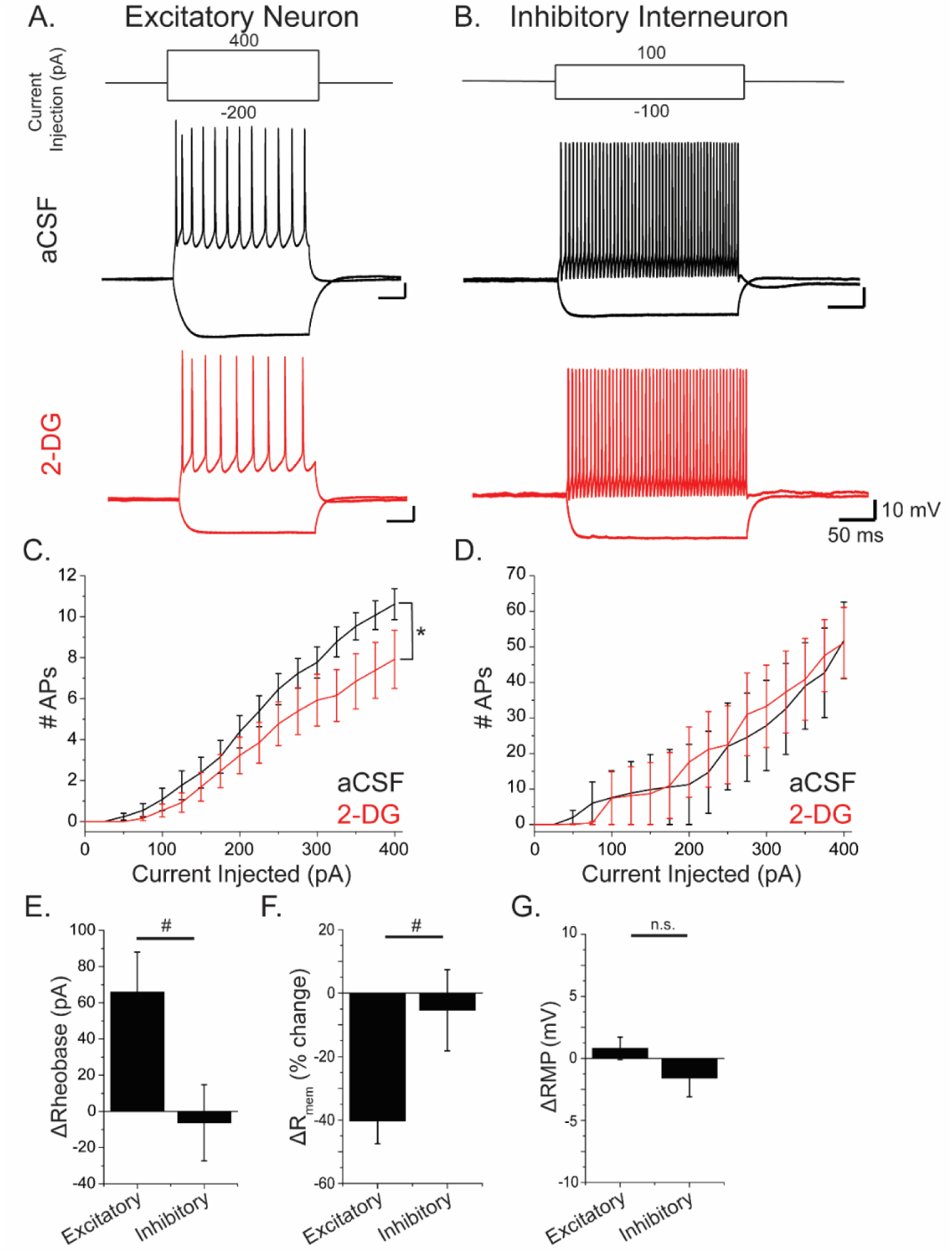
*In vitro* 2-DG treatment decreases the intrinsic excitability of excitatory pyramidal neurons. **A-B**. Representative traces following current injection into Layer V cortical excitatory pyramidal neurons **(A)** or interneurons **(B)** before (black) or after (red) treating the cortical slice with 2-DG for 10 minutes. **C.** Input-output curves from excitatory pyramidal neurons. **D.** Input-output curves from inhibitory interneurons. **E.** ΔRheobase (current injection required to fire the first action potential in 2-DG versus baseline) in excitatory neurons and interneurons. **F.** % change in membrane resistance of excitatory neurons and interneurons following 2-DG treatment. **G.** Change in resting membrane potential (RMP) in excitatory neurons and interneurons in 2-DG. (Error bar = SEM; LMM: *t > ±1.96, effect: interaction between current injection and 2-DG; # t > ±1.96, effect: cell type; n.s. not significant)

To control for reduced 2 mM glucose in 2-DG-aCSF, these experiments were repeated in low glucose aCSF (LG-aCSF: 2 mM glucose, 8 mM sucrose (a metabolically inert sugar)). LG-aCSF had no effects on AP firing, input-output curves, or membrane resistance in either excitatory neurons or inhibitory interneurons (Figure S1). This demonstrates that simply lowering glucose to 2 mM is not sufficient to alter neuronal excitability. Together, these experiments show that inhibition of glycolysis has cell type-specific effects on neuronal excitability.

### 2-DG ameliorates CCI-induced increases in synaptic excitation onto GABAergic interneurons

Acutely after TBI, glycolytic activity and neuronal activity are aberrantly increased (*35*). We hypothesized that TBI results in increased synaptic excitation of GABAergic interneurons in the peri-injury cortex acutely after injury. If so, this hyperexcitation may contribute to later cell death or dysfunction of PV (and other GABAergic) interneurons. To test this hypothesis, we utilized controlled cortical impact (CCI) to model focal, contusional TBI in adult male mice. Briefly, a craniectomy was made over left sensorimotor cortex and an impact was delivered to generate a moderate-to-severe injury (3 mm impactor probe diameter, 3.5 m/s velocity, 400 ms dwell time, 1 mm impact depth) (*44*). Acute cortical slices were prepared from sham- and CCI-injured brains 3 days following injury and whole-cell recordings were made from fast-spiking GABAergic interneurons within 200 µm of the lateral edge of the CCI lesion. Neurons were voltage-clamped at −70 mV to isolate spontaneous excitatory postsynaptic currents (sEPSCs). Consistent with our hypothesis, there was an increase in the mean frequency of sEPSCs after CCI compared to sham (Figure 2B; LMM to examine effect of CCI; N = 17 Sham and 20 CCI cells from 4 animals/group; t = 2.493). CCI also resulted in a significant leftward shift in the sEPSC inter-event interval cumulative distribution (Figure 2C; 2-sample K-S test, sham vs. CCI, p = 6.84E-11). Similarly, the mean sEPSC amplitude was increased, although this did not reach statistical significance based on LMM, likely due to large within-group variability in the outcome (Figure 2E-F; effect size = 3.144 pA; t = 1.773). There was a significant rightward shift the sEPSC amplitude cumulative distribution (Figure 2G; 2-sample K-S test, sham vs. CCI, p = 1.14E-4).

**Figure 2:**
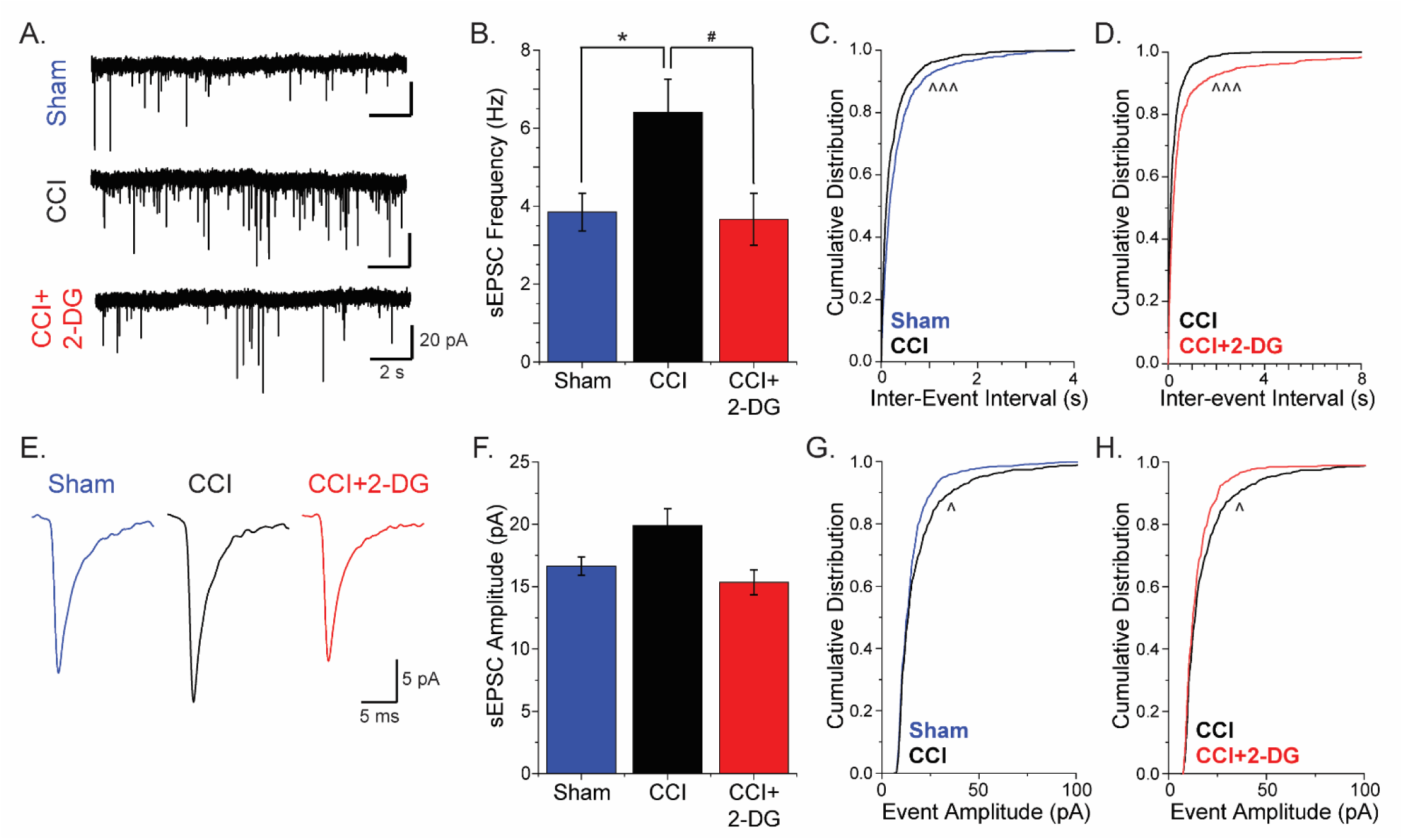
*In vitro* 2-DG attenuates CCI-induced increases of excitatory synaptic activity onto interneurons 3 days after injury. **A.** Representative sEPSC traces from acute cortical slices from sham animals or CCI-injured animals, and a representative trace from a CCI slice following 10 minutes of 2-DG wash-on. **B.** Mean sEPSC frequencies. **C-D.** Cumulative distributions of inter-event interval generated from 100 random events per recorded cell. **E.** Representative events from each condition, generated by averaging 100 events from a representative trace. **F.** Average event amplitude. **G-H.** Cumulative distributions of event amplitude generated from 100 randomly selected events per cell. (Error bar = SEM; LMM: *t > ±1.96, effect: CCI; #t > ±1.96, effect: interaction between CCI and 2-DG; 2-sample K-S test: ^p < 0.001; ^^^p < 1E-5)

Because glucose utilization is elevated acutely after TBI and we had previously observed 2-DG’s specific effects on excitatory (but not inhibitory) neuronal activity, we assessed whether glycolytic inhibition with 2-DG could reduce injury-induced excitation of GABAergic interneurons. When 2-DG-aCSF was applied to acute cortical slices prepared 3 days following CCI, the mean frequency of sEPSCs onto interneurons was reduced to approximately sham levels (Figure 2B; LMM, fixed effect: interaction between CCI and 2-DG; N = 20 CCI and 15 CCI+2-DG cells from 4 animals/group; t = −3.036), corresponding with a rightward shift the sEPSC inter-event interval cumulative distribution (Figure 2D; 2-sample K-S test, CCI vs. CCI+2-DG, p = 1.23E-13). Application of 2-DG to sham tissue did not significantly affect sEPSC frequency. 2-DG treatment also resulted in a leftward shift of the cumulative event amplitude following CCI (Figure 2H; 2-sample K-S test, CCI vs. CCI+2-DG, p = 8.07E-4) while comparisons of the mean amplitude by LMM did not reach statistical significance. Thus, this experiment suggests that GABAergic interneurons receive increased glycolysis-dependent excitation in the days following TBI.

### Acute 2-DG treatment attenuates epileptiform activity in cortical brain slices following CCI

Next, we tested whether 2-DG application could attenuate network-level cortical hyperexcitability, which is observable in field excitatory post-synaptic potentials (fEPSPs) as early as 2 weeks following TBI (*17*), and is robustly present by 3-5 weeks following TBI. Brain slices were prepared from mice 3-5 weeks after CCI or sham surgery and electrical stimulation was delivered to ascending cortical inputs to evoke fEPSPs. In sham animals, fEPSPs were brief and low-amplitude. Slices taken 3-5 weeks after CCI, however, exhibited stimulus-evoked fEPSPs with epileptiform activity (increased amplitude, duration, area, and coastline), consistent with network hyperexcitability ((*20*), Fig. 3A). As shown previously in other slice models of epilepsy (*40*), *in vitro* application of 2-DG markedly reduced the percentage of stimulus-evoked fEPSP traces that exhibited epileptiform activity after CCI from 90% to 18% (Figure 3B; LMM, fixed effect: interaction between CCI and 2-DG; N = 10 slices from 6 animals; t = −8.825). Additionally, the 2-DG treatment resulted in field potentials with significantly decreased area (Figure 3C; LMM, fixed effect: interaction between CCI and 2-DG; t = −3.047) and high-frequency activity as measured by fEPSP coastline (Figure 3D; LMM, fixed effect: interaction between CCI and 2-DG; t = −3.859). In brain slices from sham-injured animals, 2-DG-aCSF did not significantly change the area or coastline of the fEPSPs, or the percentage of epileptiform traces. In low glucose conditions (LG-aCSF), post-CCI epileptiform activity was not attenuated (Figure S2). These results show that *in vitro* glycolytic inhibition with 2-DG rapidly attenuates cortical network hyperexcitability 3-5 weeks after TBI.

**Figure 3:**
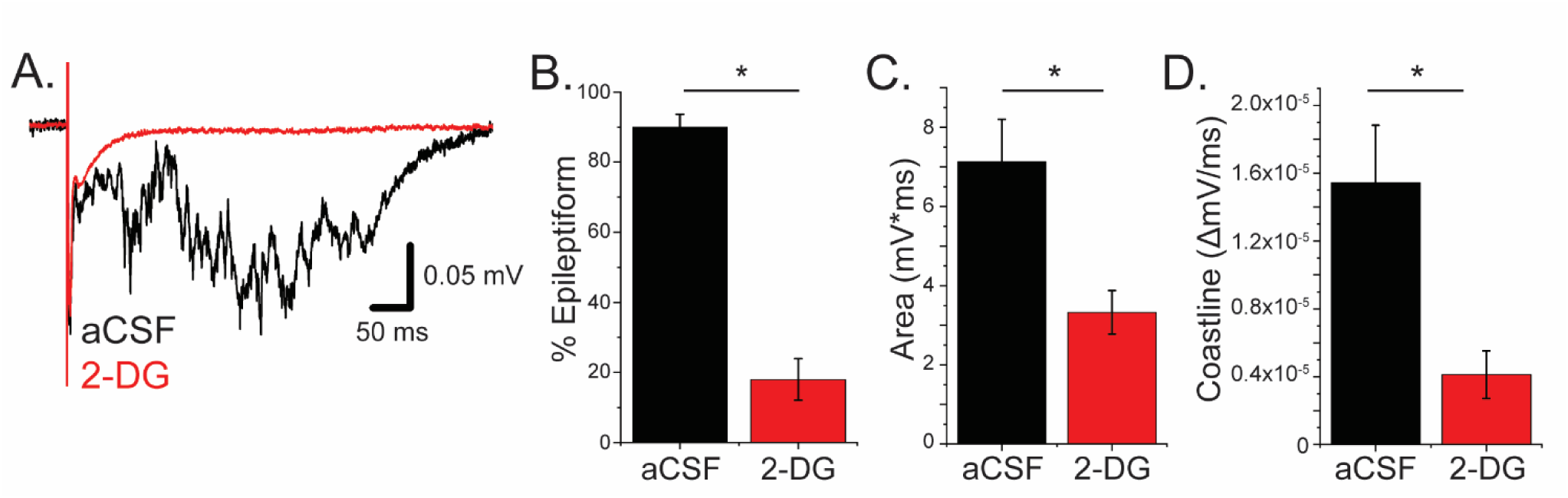
Acute 2-DG treatment decreases epileptiform activity *in vitro* following CCI. **A.** Representative stimulus-evoked field potentials in an acute cortical slice from a CCI-injured animal before (black) and after (red) local perfusion of 2-DG for 30 minutes. **B.** Percentage of traces with epileptiform activity after CCI. **C-D.** Area (C) and coastline (D) of the field potentials in baseline aCSF and 2-DG-acSF in CCI-injured slices. (Error bar = SEM; LMM: *t > ±1.96, effect: interaction between CCI and 2-DG)

### In vivo 2-DG treatment attenuates epileptiform cortical activity following CCI

We next hypothesized that reducing glycolysis *in vivo* during a critical period following TBI would reduce subsequent cortical hyperexcitability. To test this hypothesis, we treated mice *in vivo* with 2-DG for the first week after injury (250 mg/kg or vehicle intraperitoneally, once daily for seven days following CCI or sham surgery) and assessed network hyperexcitability. Acute cortical brains slices were prepared 3-5 weeks following the initial surgery (2-4 weeks following the end of 2-DG treatment), when epileptiform activity would be expected in the CCI animals (*20*). As predicted, sham-injured slices from both treatment groups had normal fEPSPs (Figure 4A-B). Consistent with previous studies, acute cortical brain slices from vehicle-treated CCI-injured animals exhibited epileptiform stimulus-evoked fEPSPs (Figure 4C). However, *in vivo* 2-DG treatment following CCI dramatically attenuated epileptiform fEPSPs (from 95% to 13%) in CCI-injured animals (Figure 4D, 4E; LMM, fixed effect: interaction between CCI and 2-DG treatment; N = 9 slices from 3 animals/group; t = −3.27). Area and coastline measurements of the evoked fEPSPs following CCI also exhibited significant attenuation with *in vivo* 2-DG treatment (Figure 4F-G; t = −3.119 and −2.102, respectively).

**Figure 4:**
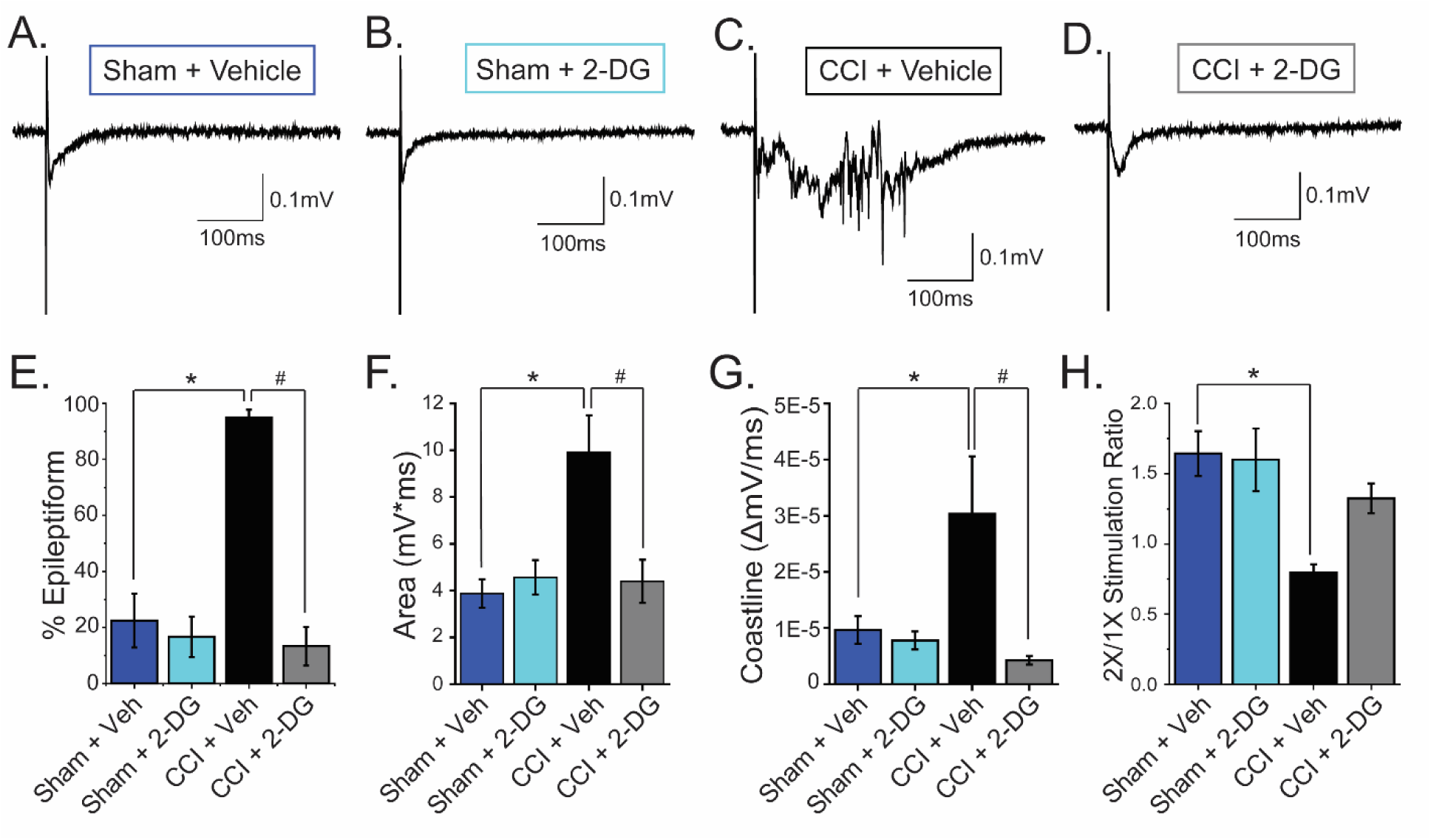
*In vivo* 2-DG treatment following CCI decreases epileptiform activity in acute cortical slices. Animals were given sham or CCI surgery, followed by 1 week of daily intraperitoneal injections of vehicle or 2-DG (250 mg/kg). **A-D.** Representative stimulus-evoked field potentials taken from acute cortical slices 3-5 weeks following sham or CCI surgery, with vehicle or 2-DG treatment. **E.** The percentage of sweeps exhibiting epileptiform activity in sham and CCI-injured animals, with or without 2-DG treatment. **F-G.** Area (F) and coastline (G) of traces taken from sham and CCI-injured animals, with or without 2-DG treatment. **H.** Input-output ratios in each treatment group. (Error bar = SEM; LMM: *t > ±1.96, effect: CCI; # t > ±1.96, effect: interaction between CCI and 2-DG)

CCI also disrupts the input/output (I/O) relationship of the cortical network. Cortical I/O responses were assessed by comparing fEPSP area evoked by 1X and 2X threshold stimulation. In the uninjured brain, cortical networks generated graded output in response to increasing stimulation. After CCI, however, cortical networks generated all-or-none responses and lacked graded output in response to stimulation (Figure 4H; LMM, fixed effect: CCI; t = −3.494). *In vivo* 2-DG treatment partially restored the I/O relationship, resulting in graded fEPSP amplitude and area in response to increasing stimulation. 2-DG treatment had a large effect size on I/O relationship (0.54 compared to the effect of CCI, −0.81) but did not reach statistical significance by LMM, likely due to the high degree of variability (Figure 4H; LMM, fixed effect: interaction of CCI and 2-DG; t = 1.568). Together, these experiments show that *in vivo* 2-DG treatment during the first week after injury attenuates network hyperexcitability and reduces long-term TBI-associated cortical network dysfunction.

### In vivo 2-DG treatment transiently affects systemic glucose metabolism, but does not affect body temperature

Since systemic metabolic changes, such as hypothermia, may have independent neuroprotective effects after TBI, we assessed the effects of *in vivo* 2-DG on whole body metabolism. After a single 2-DG injection (250 mg/kg, intraperitoneal) in fasted animals, blood glucose levels were transiently increased (Figure S3A; LMM, fixed effect: effect of 2-DG at Hour 2; N = 4 vehicle, 6 2-DG animals; t = 3.310) and β-hydroxybutyrate (a ketone body) levels were transiently decreased (Figure S3B; LMM, fixed effect: effect of 2-DG at Hour 2; t = −3.586), consistent with decreased glucose utilization and increased ketosis. These changes in blood glucose and β-hydroxybutyrate returned to baseline by 4 hours post-injection (LMM, fixed effect: effect of 2-DG at Hour 4; t = 0.010, −0.775, respectively). There was also a transient decrease in home cage locomotor activity that resolved to vehicle levels after the first 6 hours post-injection (Figure S3C; 2-sample t-test H0-6, p = 2.99E-4; H7-12, p = 0.891). Importantly, there was no change in flank temperature in 2-DG-injected animals versus vehicle-injected controls (Figure S3D; LMM, fixed effect: effect of 2-DG at Hour 2; t = 1.328), suggesting that hypothermia does not contribute to the neuroprotective effect of 2-DG.

Then, one week of daily 2-DG injections was performed in naïve animals to parallel the *in vivo* dosing regimen post-CCI. Animals were assessed while at rest, immediately before their daily 2-DG injection (~24 hours following their most recent injection). There were no changes observed in blood glucose, blood β-hydroxybutyrate, flank temperature, body weight, or daily food intake between vehicle- and 2-DG-treated animals across the entire week of treatment (Figure S4). For all experiments reported in this study, *in vivo* treatment with 2-DG did not affect animal weight or recovery from surgery. These data suggest that there were no significant adaptive metabolic changes across the week-long *in vivo* 2-DG treatment.

### In vivo 2-DG treatment prevents CCI-induced dysregulation of excitatory and inhibitory synaptic activity

TBI causes significant disruption of cortical synaptic activity including increased glutamatergic excitation and reduced GABAergic inhibition of excitatory cortical neurons (*20*), accordant with network hyperexcitability. Consistent with previous studies, CCI induced a significant increase in mean spontaneous excitatory post-synaptic current (sEPSC) frequency (Figure 5A; LMM, fixed effect: CCI; N = 22 sham and 26 CCI cells from 5 animals/group; t = 2.82), resulted in a significant leftward shift in the sEPSC inter-event interval cumulative distribution (2-sample K-S test, sham vs. CCI, p = 1.44E-46), and had no effect on sEPSC amplitude (Figure 5C). Inhibitory synaptic transmission was decreased by CCI, again consistent with previous studies. CCI decreased mean spontaneous inhibitory post-synaptic current (sIPSC) frequency (Figure 5G; LMM, fixed effect: CCI; N = 25 sham and 21 cells from 5 animals/group; t = −2.023), caused a significant rightward shift in the sIPSC inter-event interval cumulative distribution (2-sample K-S test, sham vs. CCI, p = 6.7E-13), and significantly increased mean sIPSC amplitude (Figure 5I; LMM; t = 3.528).

**Figure 5:**
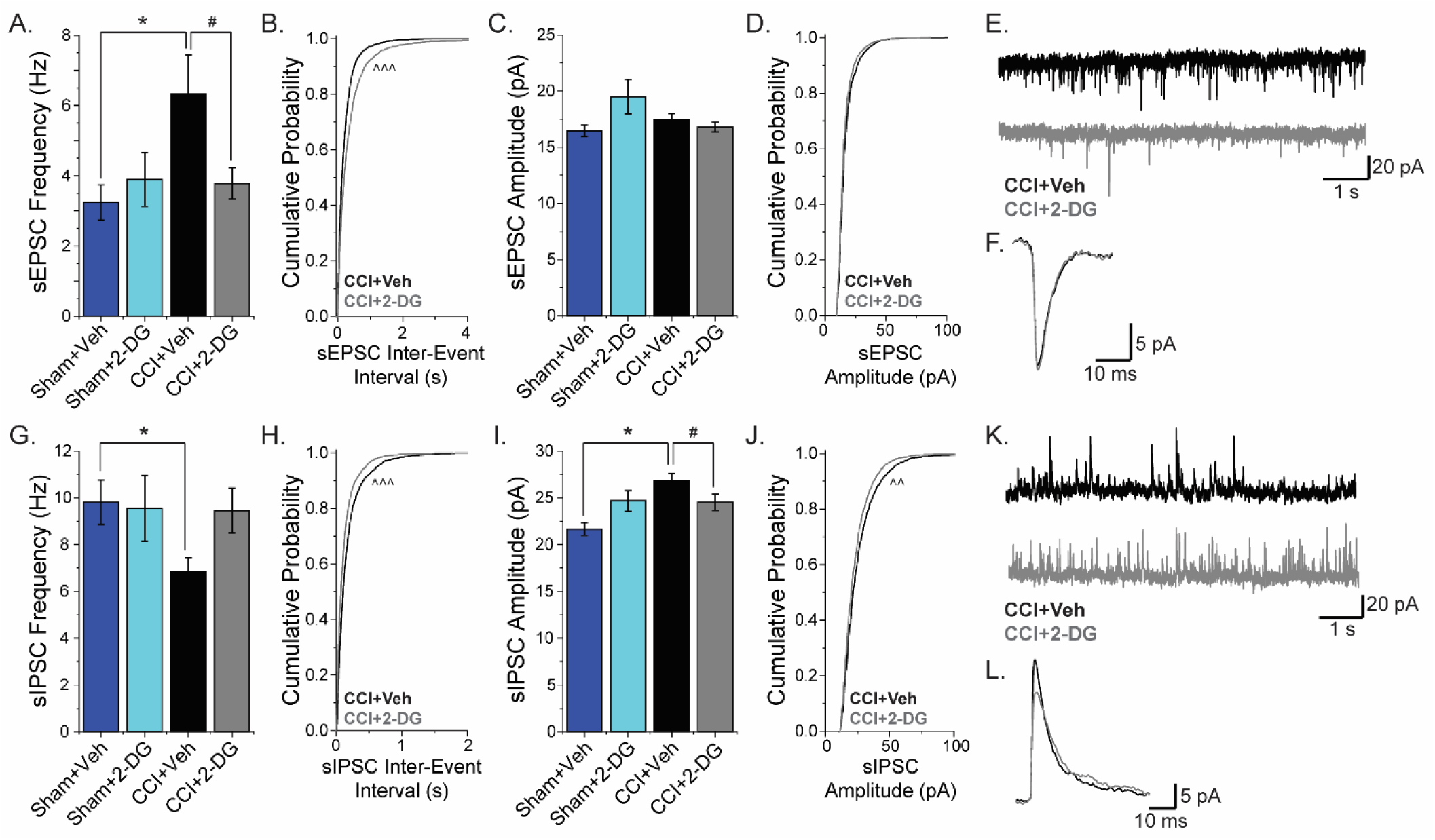
*In vivo* 2-DG treatment attenuates changes in synaptic activity after injury. **A.** Mean sEPSC frequency. **B.** Cumulative distribution of sEPSC inter-event interval. **C.** Mean sEPSC amplitude. **D.** sEPSC amplitude cumulative distribution. **E.** Representative sEPSC traces from vehicle-treated or 2DG-treated CCI brains. **F.** Representative example of sEPSC events (generated by averaging 100 events from a representative trace). **G.** Mean sIPSC frequency. **H.** Cumulative distribution of sIPSC inter-event interval. **I.** Mean sIPSC amplitude. **J.** Cumulative distribution of sIPSC amplitude. **K.** Representative sIPSC traces from vehicle-treated or 2-DG-treated CCI brains. **L.** Representative example of sIPSC events from vehicle-treated or 2-DG-treated CCI brains. (Error bar = SEM; LMM: *t > ±1.96, effect: CCI; #t > ±1.96, effect: interaction between CCI and 2-DG; 2-sample K-S test: ^^p < 1E-4; ^^^p < 1E-5)

Consistent with its effects on network hyperexcitability, *in vivo* 2-DG treatment significantly attenuated CCI-induced increases in excitation. 2-DG treatment reduced mean sEPSC frequency (Figure 5A, 5E; LMM, fixed effect: interaction between CCI and 2-DG; N = 26 CCI+Vehicle and 22 CCI+2-DG cells from 5 animals/group; t = −2.032) and caused a significant rightward shift in the sEPSC inter-event interval cumulative distribution (Figure 5B; 2-sample K-S test, p = 1.55E-20), with no effect on sEPSC amplitude (Figure 5C-D; LMM; fixed effect: interaction between CCI and 2-DG, t = −1.762). *In vivo* treatment with 2-DG also restored inhibitory synaptic transmission after CCI. 2-DG had a large, but not significant, effect on mean sIPSC frequency (Figure 5G; LMM, fixed effect: interaction between CCI and 2-DG; N = 21 CCI+Vehicle and 21 CCI+2-DG cells from 5 animals/group; t = 1.371), caused a significant leftward shift in the sIPSC inter-event interval cumulative distribution (Figure 5H; 2-sample K-S test, p = 7.03E-10), significantly reduced mean sIPSC amplitude (Figure 5I; LMM, fixed effect: interaction between CCI and 2-DG; t = −2.483), and caused a leftward shift in the cumulative distribution of sIPSC amplitude (Figure 5J; 2-sample K-S test, p = 1.46E-5). Interestingly, we noted a difference in the distribution of mean sIPSC frequency following CCI; in cells recorded after sham injury, there was consistently a population of neurons (5 out of 25 cells in the Sham+Vehicle group) that received sIPSCs at a high rate (>12 Hz). This population was absent after CCI (0 out of 21 cells in CCI+Vehicle) but was present after *in vivo* 2-DG treatment (7 out of 21 cells in CCI+2-DG), suggesting that CCI causes a loss of robust network inhibition that is prevented by 2-DG treatment. Together, these data demonstrate that *in vivo* 2-DG treatment reverses deficits in both excitatory and inhibitory neurotransmission caused by CCI.

### In vivo 2-DG treatment attenuates the loss of parvalbumin-expressing interneurons following CCI

Because 2-DG treatment attenuated the development of epileptiform activity and prevented decreases in sIPSCs after CCI, we hypothesized that it may also prevent the loss of parvalbumin-expressing interneurons that had been previously reported (*20*). An additional question we wished to address was whether CCI caused a loss of PV cells, or merely a loss of PV protein. To test our hypothesis and address the loss of PV cells versus protein, we used a combination of genetic and immunohistochemical approaches. *PV*^*Cre*^ mice, in which Cre-recombinase is expressed under the control of the parvalbumin promoter, were crossed with *Ai9* reporter mice, producing *PV*^*Cre*^/*Ai9* mice which express tdTomato (tdT) in cells containing Cre. This provides a genetic label of PV cells, even if the PV protein itself is lost.

*PV*^*Cre*^/*Ai9* mice underwent CCI or sham surgeries and were treated *in vivo* with 2-DG or vehicle as described above. Brains were prepared for immunohistochemical analysis 3-5 weeks post-CCI. First, lesion volume was approximated across serial sections between CCI animals treated with vehicle or 2-DG. *In vivo* 2-DG treatment had no significant effect on cavitary lesion volume (2-sample t-test; N = 5 animals/group, p = 0.557). Then, the densities of PV-immunolabeled and tdT-labeled neurons were quantified in five 100-μm-wide rectangular regions of interest (ROIs) in the cortex lateral to the site of CCI injury (or in isotopic cortex in sham animals). Consistent with previous studies (*20*), the density of PV-immunolabeled cells was significantly reduced in the cortex of vehicle-treated CCI animals relative to sham (Figure 6A-D, 6G; LMM with Type III ANOVA – reports global effects across all ROIs; N = 3 slices/animal from 5-7 animals/group; effect: CCI across all ROIs, p = 5.83E-15). PV+ cell density was significantly reduced in the first 400 µm adjacent to the lesion (Figure 6G; LMM, fixed effect: interaction between CCI and ROI1-4; t < −1.96). A similar result was seen after counting of tdT-labeled neurons (Fig. 6H; LMM with Type III ANOVA; effect: CCI across all ROIs, p = 1.48E-11) although the decrease was significant only in the first 200 µm lateral to the lesion (LMM, fixed effect: interaction between CCI and ROI1-2; t < −1.96). These results show that CCI causes both a loss of PV cells and a decrease in PV immunoreactivity adjacent to the lesion.

**Figure 6:**
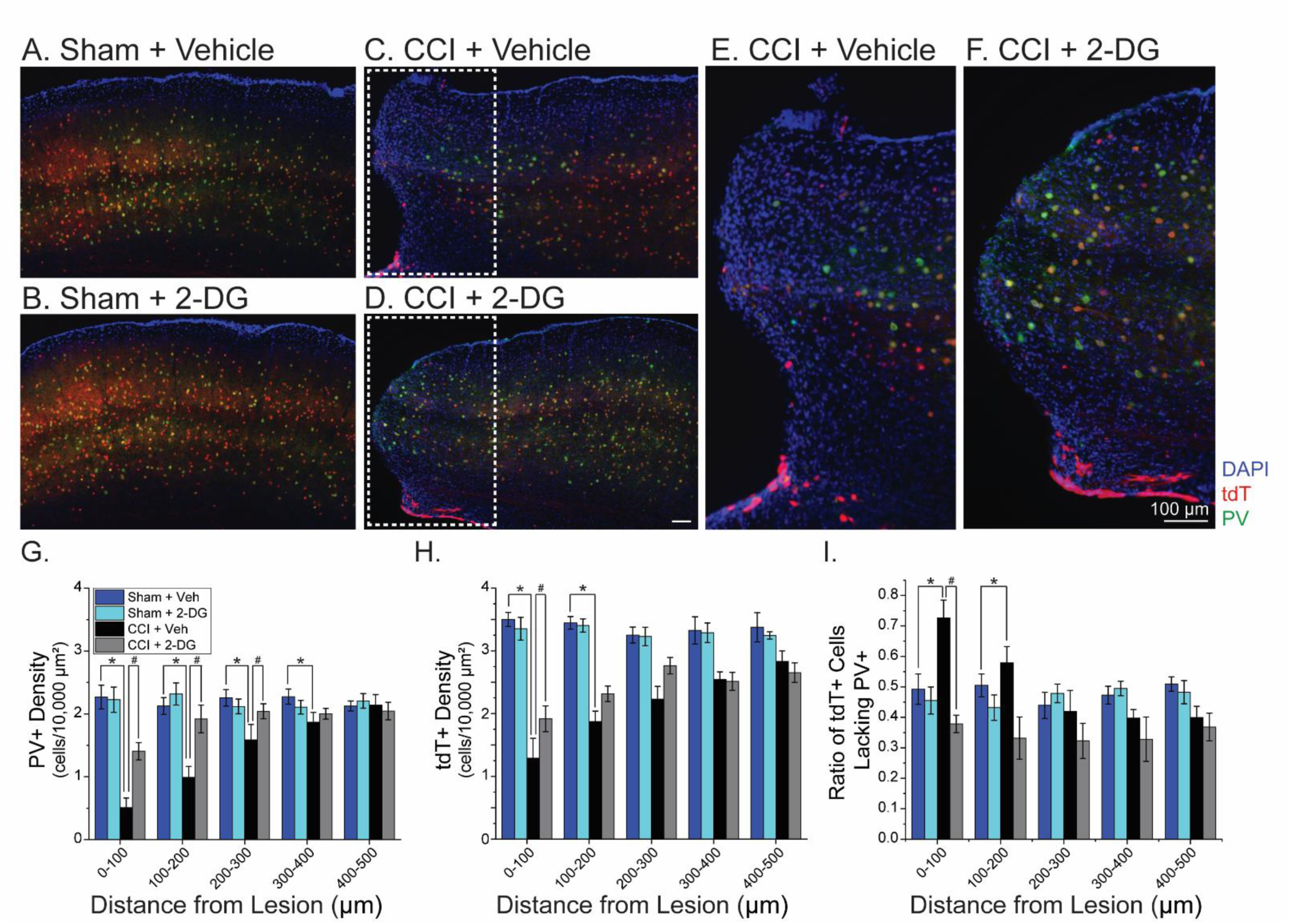
*In vivo* 2-DG treatment attenuates the decrease in PV+ cells in the perilesional cortex. **A-D.** Representative images with PV (green), tdT (red – in *PV*^*Cre*^ cells), and DAPI (blue) in animals from the different treatment groups, collected 3-5 weeks after sham or CCI surgery. **E-F.** Zoom-in of perilesional area showing loss of colocalized PV+tdT+ cells in the perilesional area after CCI. **G.** Quantification of PV+ cell density in five regions lateral to the CCI cavitary lesion. **H.** Quantification of cells identified with tdT+ in *PV*^*Cre*^/*Ai9* reporter mice. **I.** The density of tdT+ cells not colocalized with PV immunolabel was divided by the total density of tdT+ cells in each ROI to calculate the ratio reported. (Error bar = SEM; LMM: *t > ±1.96, effect: CCI in each ROI; #t > ±1.96, effect: interaction of CCI and 2-DG in each ROI)

*In vivo* 2-DG treatment attenuated the loss of PV+ neurons relative to vehicle-treated CCI animals (Figure 6F-G; LMM with Type III ANOVA; interaction of CCI and 2-DG treatment across all ROIs; p = 0.003). 2-DG prevented PV+ cell loss in the first 300 µm adjacent to the lesion (LMM, fixed effect: interaction between CCI, 2-DG, and ROI1-3; t > 1.96). 2-DG did not globally reduce the loss of tdT+ neurons (LMM with Type III ANOVA; interaction of CCI and 2-DG treatment across all ROIs; p = 0.112), although a significant increase in tdT+ neurons was observed in the 100 µm closest to the site of injury (Figure 6H; LMM; fixed effect: interaction between CCI, 2-DG, and ROI1, t > 1.96), suggesting a focal effect of 2-DG. These results indicate that 2-DG broadly attenuates the loss of PV expression after CCI and focally reduces the loss PV (tdT+) neurons.

In order to further examine the effects of CCI and 2-DG on PV cell loss, versus changes in PV protein expression, we assessed the proportion of tdT-labeled neurons that were immune-positive for PV. In sham-injured animals, approximately 50% of tdT-labeled neurons were PV immuno-positive. The abundance of tdT-labeled neurons which lacked PV immunolabeling is likely due to a combination of PV expression below the detection limit of an immunolabeling approach, transient expression of PV during development, and non-specific Cre expression. After CCI, a significantly higher proportion of tdT-labeled neurons lacked PV immunoreactivity (Figure 6I; LMM with Type III ANOVA; effect: CCI across all ROIs, p = 2.78E-6) with the effect being most prominent in the first 200 µm adjacent to the lesion (LMM, fixed effect: interaction between CCI and ROI1-2; t > 1.96). This suggests that a sub-set of PV (tdT+) interneurons remain after CCI but lose their expression of PV protein. Because PV is a calcium binding protein that is necessary for physiologic interneuron AP firing, loss of PV expression is consistent with decreased inhibitory function and cortical hyperexcitability (*50, 51*). *In vivo* treatment with 2-DG restored the ratio of PV (tdT+) cells expressing PV to levels seen in sham animals (Figure 6I; LMM with Type III ANOVA; effect: interaction of CCI and 2-DG across all ROIs, p = 1.37E-3). Together, these results show that CCI causes a loss of PV (tdT+) interneurons and reduces PV expression in remaining PV interneurons. *In vivo* 2-DG treatment broadly increases PV expression in surviving PV interneurons, consistent with enhanced inhibitory function.

## Discussion

2-DG, a hexokinase inhibitor that reduces glycolytic activity, has been explored for its potential neuroprotective, anti-convulsant, and anti-epileptogenic properties, but the mechanisms of its rapid actions on neuronal function are unknown. Here, we demonstrate that 2-DG treatment attenuates excitatory, but not inhibitory, cortical neuron excitability. This cell type-specific coupling of neuronal excitability to metabolic activity provides a mechanism which engenders 2-DG with acute anticonvulsant properties. Further, we show that 2-DG has therapeutic potential to treat traumatic brain injury (TBI). While glycolysis is known to be abnormally elevated acutely following brain injury (*52*), it has been previously unknown whether early metabolic disruptions contribute to chronic cortical hyperexcitability. Using the CCI model of TBI combined with electrophysiological approaches, we identify aberrant excitation onto inhibitory interneurons 3 days after injury that can be attenuated by acute 2-DG treatment. This provides novel insight into acute synaptic dysfunction following TBI, implicates glycolysis as possibly mediating the hyperexcitation of inhibitory interneurons in this setting, and suggests that 2-DG treatment may reduce post-traumatic inhibitory network dysfunction.

In addition to early changes in inhibitory synaptic activity, the loss of cortical interneurons after TBI leads to long-term synaptic dysfunction and cortical hyperexcitability. We suspected that reducing hyperexcitation of GABAergic interneurons after TBI would preserve inhibitory network function following CCI. To test this hypothesis, we treated animals *in vivo* with 2-DG during the first week after TBI. We found that this approach prevented epileptiform network activity, synaptic dysfunction, and parvalbumin-positive inhibitory interneuron loss associated with CCI. Our results demonstrate that 2-DG treatment reduces TBI-related pathologies and that modulation of metabolism in the injured brain may have theraperutic potential to improve patient outcomes.

From a translational standpoint, the therapeutic rationale to reduce glycolysis after TBI is well-founded on both preclinical and clinical literature (*35*). Using PET scanning, studies report that a majority of patients exhibit increased cerebral glucose utilization following severe TBI ((*52, 53*)) that later develops into a chronic, hypometabolic state (*54–56*). Additionally, patients with aberrant glucose utilization and network activity during the first week post-TBI have worsened outcomes 6 months later (*57*). Preclinical work from multiple animal models of TBI shows early increases in glucose utilization ((*33, 58–61*)) which are activity-dependent (*34*). Consistent with focal hyper-glycolysis and network dysfunction, our previous studies show heightened glutamate signaling in the peri-injury cortex and parvalbumin-interneuron loss also regionally confined to this area. Our approach to utilize 2-DG after TBI parallels 2-DG’s use to limit tumor growth in cancer, where neoplastic cells exhibit increased glycolytic activity via the Warburg effect (*62*). In our case, 2-DG may specifically target regions and cells exhibiting hyper-glycolysis to attenuate aberrant excitatory neuron activity and minimize long-term pathological changes in the cortex.

Supporting 2-DG’s translational potential, we did not find evidence of chronic changes in animal health or systemic metabolism over the course of 2-DG treatment. Transient changes in metabolism and locomotion occur in the first few hours after 2-DG injection, as previously reported (*63*). Long-term exposure to 2-DG, however, has been previously associated with cardiotoxicity (*64, 65*), underscoring that short-term use as a disease-modifying agent (as in our approach) is preferable over chronic use as an anticonvulsant. Finally, there is abundant evidence supporting the use of the ketogenic diet (which results in decreased glycolysis and increased ketosis) as a strategy to manipulate neural network activity in animals and humans (reviewed in (*66–72*)). While dietary approaches and glycolytic inhibition have shown efficacy in controlling seizure activity, manipulations of glucose metabolism have not yet been harnessed effectively in the context of TBI.

A particularly important finding of our study is the cell type-specific coupling of metabolism to neuronal excitability. Our results are concordant with published studies showing that GABAergic interneurons have unique energetic requirements and are known to be enriched with mitochondria for energy production via oxidative phosphorylation (reviewed in (*73*)). Our findings suggest that glutamatergic pyramidal neurons rely on glycolysis to maintain their intrinsic excitability. While glycolysis is traditionally thought to occur mainly in astrocytes (astrocyte-neuron lactate shuttle model (*74*)), there is significant evidence that glycolysis also occurs in neurons. Multiple glycolytic enzymes are enriched in neurons relative to astrocytes (*75, 76*) and recent work implicated increased neuronal glycolysis in conditions of heightened energy need (*77, 78*). Thus, increased energetic requirements, glycolytic flux, and cell-type specific neuronal activity may all be interdependent during the acute period following TBI, and deserve further investigation. A deeper understanding of metabolic demands following TBI and of how different CNS cells utilize energy will aid in the development of metabolic approaches to manipulate neural network activity with greater precision.

In summary, we report that *in vitro* and *in vivo* 2-DG treatment attenuates pathologic cellular and network changes following CCI in mice. A caveat of the studies presented is that they do not specifically examine the degree of glycolytic inhibition invoked by our *in vivo* 2-DG treatment paradigm, and whether glycolysis is primarily targeted in neurons, astrocytes, immune cells, or other cells in the post-TBI cortex. Future work is needed to investigate the use of glycolytic inhibition to prevent injury pathology, including in diffuse or concussive mechanisms of TBI, in larger animal models, and in pilot clinical trials. Additionally, studies are needed to explore whether reducing cortical hyperexcitability is sufficient to ameliorate patient-oriented outcomes including mortality, seizure frequency/severity in post-traumatic epilepsy, and cognitive/behavioral outcomes. Finally, improving our understanding of the specific energy demands of the neuronal activity, hemodynamic response, and inflammation that occur after TBI will enable us to harness metabolic manipulations to reduce brain injury-associated pathologies. The findings of this study support continued investigation into the use of glycolytic inhibition as a therapeutic approach to prevent post-TBI complications.

## Materials and Methods

### Study Design

#### Basic Design

All experiments were hypothesis-driven, rigorously designed, and performed in a controlled laboratory environment. Sample size was set based on power analysis calculations using GPower to provide power > 0.80 for α = 0.05 for most experiments. We also utilized standards in the field, i.e. ≥3 animals per group, to ensure proper statistical power.

#### Randomization and blinding

Mice were randomized to group within each experiment (sham vs. CCI; vehicle vs. 2-DG). The studies were blinded at the level of data analysis. Data was numerically encoded and analyzed by a trained experimenter blinded to condition. After initial analysis was finalized, the blinding was lifted to compare data across groups.

#### Exclusion criteria and statistical approach

Each experiment had one primary endpoint based on the experimental design. We did not define rules for stopping data collection outside of humane endpoints based on animal protocols. Data was excluded based on quality control of whole-cell recordings (recordings were excluded when access resistance changed more than 20% during an experiment). Outliers were removed in the context of fEPSP recordings due to inherent variability in the responses evoked with this technique (Figures 3-4). Outliers were defined as slices with response area > 2 standard deviations from the group mean and were excluded from the subsequent analysis. The number of experimental replicates is included in the text. The number of individual animals, cells, or slices were factored into our analysis using a linear mixed model approach. This approach includes terms to account for inter- and intra-animal variability.

### Animals

All animal procedures were performed in accordance with the Tufts University School of Medicine’s Institutional Animal Care and Use Committee. All experiments were performed on adult mice. C57/BL6 were obtained from Jackson Labs (Strain #000664), Charles River Laboratories (Strain #027), or bred in-house. *PV*^*Cre*^ (Stock # 017320), *Ai9* (Stock # 07909), and *G42* (Stock # 007677) mice were obtained from Jackson Laboratories. Animals were kept on a standard 12-hour light cycle and fed *ad libitum* with regular chow diet and water. Male mice were used for all experiments, unless noted.

### Controlled cortical impact

Traumatic brain injury was modeled with controlled cortical impact (CCI), as previously described (*20, 79*). Briefly, 10-14-week-old male mice were anesthetized using inhaled isoflurane in oxygen (4% for induction, 2% for maintenance). Following placement in a Kopf stereotaxic frame, the surgical area was sterilized and a vertical, midline skin incision was made. A 5 mm craniectomy was performed lateral to midline, between bregma and lambda (over the left somatosensory cortex) and the skull flap was removed. The surgical field was flushed with sterile saline throughout the procedure to cool the surgical area during the craniectomy drilling. Impact was performed with a Leica Benchmark Stereotaxic Impactor, using a 3 mm diameter piston, 3.5 m/s velocity, 400 ms dwell time, and 1 mm depth. After the CCI procedure, sutures were used to close the incision. The bone flap was not replaced in order to accommodate swelling after the procedure and to prevent pressure-induced damage to the injury site. Sham animals received the anesthesia and craniotomy drilling, but did not receive the CCI. Animals were singly housed following surgery until time of sacrifice.

### In vivo 2-DG treatment

Animals were injected intraperitoneally (I.P.) with 250 mg/kg 2-DG dissolved in sterile saline to a final injection volume of 200 μL per dose. Vehicle injections consisted of 200 μL of sterile saline per dose. For the *in vivo* 2-DG treatment paradigm, the first dose was administered ~20 minutes after CCI or sham surgery, and the doses were continued daily for 7 days at the same time of day. Animal weight was recorded daily to monitor animal health.

### Acute brain slice preparation

Brain slices were prepared as previously described (*20, 80*). Briefly, mice were anesthetized in an isoflurane chamber and then decapitated by guillotine. Brains were rapidly removed and placed in chilled slicing solution (in mM: 234 sucrose, 11 glucose, 24 NaHCO_3_, 2.5 KCl, 1.25 NaH_2_PO_4_, 10 MgSO_4_, 0.5 CaCl_2_) equilibrated with 95% O_2_/5% CO_2_. The brain was then glued to the slicing stage of a Leica VT1200S vibratome and the slicing chamber was filled with chilled slicing solution, again equilibrated with 95% O_2_/5% CO_2_. Coronal slices (300 or 400 μm-thick) were taken at 0.05 mm/s and hemisected. The slices were placed in a chamber filled with artificial cerebrospinal fluid (aCSF, in mM: 126 NaCl, 26 NaHCO_3_, 3 KCl, 1.25 NaH_2_PO_4_, 2 MgCl_2_, 2 CaCl_2_, 10 glucose) equilibrated with 95% O_2_/5% CO_2_. The chamber with the acute cortical slices was incubated in a water bath at 34°C for one hour, and then stored at room temperature until recording.

### In vitro treatment of acute cortical slices with 2-DG

For 2-DG wash-on experiments, 2-DG-aCSF solution was prepared by adding 8 mM 2-DG and removing 8 mM glucose from the standard aCSF recipe (to maintain solution osmolarity, for a final concentration of 2 mM glucose, 8 mM 2-DG). Baseline field or whole-cell recordings were obtained, and then the regular aCSF perfusate was replaced with 2-DG-aCSF for the remaining recordings. For low glucose wash-on experiments, aCSF containing 2 mM glucose and 8 mM sucrose (an inert sugar) was prepared (LG-aCSF), and LG-aCSF was washed on following baseline field or whole-cell recording collection.

### Whole-cell recordings

Acute cortical slices were placed in a submersion recording chamber of an Olympus BX51 microscope with continuous perfusion of oxygenated aCSF at 32°C. Layer V pyramidal neurons were visually identified with infrared differential interference contrast microscopy, and fast-spiking parvalbumin-positive cortical interneurons were identified based on GFP expression in *G42* mice (Jax Stock #: 007677). Whole-cell recording mode was achieved with a borosilicate glass electrode (3-5 MΩ) filled with internal solution optimized for each experiment (described below). Access resistance, membrane resistance, and capacitance were monitored throughout each experiment. Cells with >20% change in access resistance during the experiment were excluded from analysis. Data were collected using an Axon Multiclamp 700B amplifier, Digidata 1440A digitizer, and pClamp software.

For recordings of synaptic activity, the internal utilized was, in mM: 140 CsMs, 10 HEPES, 5 NaCl, 0.2 EGTA, 5 QX314, 1.8 MgATP, 0.3 NaGTP, pH = 7.25, mOsm = 290. The recording electrode was placed within 200 µm of the lateral edge of the cavitary cortical lesion. Spontaneous EPSCs and IPSCs were recorded by voltage-clamping the cell at −70 and 0 mV, respectively, for 2 minutes. Synaptic activity analysis was performed using MiniAnalysis (SynaptoSoft) with the experimenter blinded to experimental group.

For current injection experiments, the internal utilized was, in mM: 120 KGluconate, 11 KCl, 10 HEPES, 10 EGTA, 2 MgCl_2_, 2 MgATP, 0.3 NaGTP, pH = 7.3, mOsm = 290. The perfused aCSF also included DNQX (20 µM), CPP (10 µM), and GABAzine (10 µM) to block synaptic activity. Current injection steps were applied to each cell (ranging from −200 → 400 pA in 25 pA steps) and cell type was confirmed using AP shape and firing rate.

### Field recordings

Brain slices were placed in an interface chamber perfused with 34°C, oxygenated aCSF at a rate of 2 mL/min. A tungsten stimulating electrode was placed at the Layer VI-white matter boundary to stimulate ascending white matter tracts. A borosilicate glass micropipette (pulled to a resistance of ~1 MΩ) was filled with aCSF and placed in the corresponding area of Layer V cortex directly above the stimulating electrode. Recordings were performed in the perilesional cortex (within 100 μm of the injury site) or in isotopic cortex in sham animals. Electrical stimulus (8-25 μA, 100 μs pulse length) was delivered using a stimulus isolator (World Precision Instruments) at 30 second intervals. The signal was amplified with a Multiclamp 700A amplifier, digitized with a Digidata 1322A digitizer (sampling rate = 20 kHz), and recorded with pClamp software (Molecular Devices). Threshold stimulus intensity was defined as the minimal required stimulus to obtain a detectable field response (≥ 0.05 mV). “2X threshold” was defined by doubling the threshold stimulus intensity for a given slice.

### Analysis of field recordings

Evoked field potentials were analyzed using pClamp software and MATLAB. First, traces were adjusted by subtracting the baseline (the average amplitude directly before the stimulus), imposing a low-pass filter (Bessel 8-pole, 1000 Hz cutoff), and filtering for electrical 60 Hz noise. The area of the field potential was measured by integrating the charge transfer in the first 250 ms following the stimulus. Coastline measurements describe the relative amount of high-frequency activity in the field response and were measured by summing the distance between each point taken every 50 μs over a 250 ms time window following the stimulus. The percentage of epileptiform sweeps for a given slice was determined by counting the number of field potentials that exhibited epileptiform activity (prolonged, high frequency, and high amplitude) out of 10 total sweeps (over a 5 minute period).

### Immunohistochemistry

Animals were transcardially perfused with 100 mL of phosphate-buffered saline (PBS) followed by 100 mL of 4% paraformaldehyde (PFA) in 0.4 M phosphate buffer (PB). Brains were dissected out, placed in 4% PFA in PB overnight, and then moved to a 30% sucrose solution for 3 days. Forty μm thick coronal slices were taken on a Thermo Fisher Microm HM 525 cryostat. Each brain was serial sectioned, so that each collection well contained a representative set of coronal sections throughout the entire CCI lesion, with each section 400 µm apart from the next. Slices were washed with PBS and then incubated in blocking buffer (10% normal goat serum (NGS), 5% bovine serum albumin (BSA) in PBS with 0.2% Triton X-100 (PBS-T)) at room temperature for 1 hour. The slices were incubated overnight at 4°C with parvalbumin (1:1000, Swant) primary antibody in PBS-T with 5% NGS/1% BSA. Then slices were washed with PBS and incubated with secondary antibody (goat anti-mouse 488 or goat anti-rabbit Cy3, 1:500, Jackson Immuno) in PBS-T with 5% NGS/1% BSA for 2 hours at room temperature. Finally, slices were rinsed with PBS and mounted, in anatomical order, onto slides using Vectashield with DAPI. Imaging was performed using a Keyence epifluorescence microscope.

### Analysis of imaging data

Image analysis was performed using Fiji/ImageJ software. 5 regions of interest (ROIs) were drawn lateral to the edge of the cavitary CCI lesion, with each ROI 100 µm wide. PV+ and tdTomato (tdT)+ cells were counted independently, by an experimenter blinded to treatment group. The cell counts were then divided by the area of the selected region to find the cell density. The number of PV+tdT+ colocalized cells was determined by comparing the locations of cell bodies selected in the individual PV+ and tdT+ counting phases. Three sections from each brain (representing the same anatomical regions across all subjects) were quantified and averaged to obtain a single cell density value per cell type in each ROI in each animal.

Lesion volume was evaluated in >10 serial sections from each animal, mounted and imaged in anatomical order. A blinded experimenter traced the size of the cavitary lesion in each section and the area of missing tissue was recorded. Each area was multiplied by 400 µm (the distance between the serial sections) to estimate the lesion volume.

### Metabolic studies

Blood glucose was measured using a OneTouch glucose meter and blood β-hydroxybutyrate was measured using a PrecisionXtra Ketone Monitor with blood taken from the tail. Flank temperature was measured using a subcutaneous temperature probe (BioMedic Data Systems Implantable Electronic ID Transponders IPTT-300). Food intake was measured by subtracting the final weight from the initial weight of the chow during the indicated study period. Home cage locomotor activity was measured utilizing a photobeam-break system (Tufts Behavior Core). Animals were fasted for 16-18 hours before acute measurements following 2-DG injection (Figure S3).

### Statistical analysis

Most experiments used a 2 × 2 factorial design. To assay the statistical significance of each factor (fixed effect) and the interaction between factors, we performed linear mixed model (LMM) analysis (see Table S1 and Supplementary Methods, (*81*)). For comparison of cumulative distributions of inter-event interval or amplitude of synaptic events, we utilized a 2-sample Kolmorogov-Smirnov test.

### Drugs and reagents

Salts were obtained from Sigma-Aldrich or Fisher Scientific. Glucose was obtained from Fisher Scientific. 2-deoxy-D-glucose and DNQX were obtained from Sigma-Aldrich. CPP was obtained from Abcam. GABAzine was obtained from Tocris.

## Author Contributions

CD and JK conceived of and designed experiments examining 2-DG’s effects on neuronal excitability and synaptic activity. CD, DC, and JK conceived of and designed experiments examining 2-DG’s effects on epileptiform activity and interneuron density. CD, JK, CL, and DK conceived of and designed experiments examining effects of 2-DG on metabolism. JK performed experiments and analyzed the data for Figures 1, 2, 5, 6, S1, and S2. DC performed experiments for Figures 3-4; DC and JK analyzed the data for Figures 3-4. JK and CL performed experiments and analyzed the data for Figures S3-S4. FN performed all LMM statistical analysis (Tables S1-S2). JK and CD wrote first draft of manuscript; comments from all authors were incorporated into the final draft.

## Acknowledgments

We would like to thank Dr. Christian Bjorbaek for his helpful feedback on the manuscript.

## Supplementary Methods

### Linear mixed model analysis

For most of the statistical analysis, we utilized linear mixed model (LMM) analysis. This approach estimates the effect size of each factor while accounting for intra- and inter-animal variability. LMMs were fitted with random intercepts to assess for the correlation between repeated measurements on the same mouse, and experiment-specific effects were analyzed for statistical significance. t values > 1.96 and < −1.96 were considered to be statistically significant and corresponded to 95% confidence intervals that did not cross zero. Each LMM examined both main fixed effects and interactions between the effects.

Experiments with a traditional 2 × 2 factorial design (including those for Figures 3-5) used LMM to examine fixed effects of CCI, 2-DG treatment, and the interaction between these two effects. To compare input-output curves in Figure 1, the fixed effects were current injection, 2-DG treatment, and the interaction between the two effects. LMM was also utilized to compare excitatory and inhibitory neurons in Figure 1, by examining cell type as the fixed effect.

In Figure 2, the statistical approach also paralleled the experimental design. The primary experimental question was whether synaptic activity onto interneurons was altered acutely after CCI (as this had not been shown previously); thus “Stage 1” of the statistical approach was to examine CCI as the solitary fixed effect. Then, in “Stage 2” of the experiment and statistical analysis, we examined the effects of *in vitro* 2-DG wash-on to slices taken from either CCI or sham animals (and thus used both CCI and 2-DG as fixed effects).

In Figure 6, there was another fixed effect (ROI), introduced as a categorical variable. We performed LMM with fixed effects of CCI, 2-DG, ROI, and interactions between each of these. Comparisons were made with ROI5 (the furthest region from the CCI lesion) as the reference point. To further interpret these complex results, we utilized LMM with the Type III analysis of variance (ANOVA) test with Satterthwaite’s method to assess the global significance of “ROI” as a single factor instead of each ROI as an independent factor relative to ROI5. Thus, the Type III ANOVA reports global effects across all ROIs.

### Cumulative distribution generation

Cumulative distributions were generated by randomly selecting 100 events from each recording to ensure that data from more active cells were not more heavily weighted than data from cells with fewer events. Random event selection was accomplished using a custom-written MATLAB script. Within each treatment group, randomly selected events from each cell were pooled to generate a single cumulative distribution, and distributions were compared using a 2-sample Kolmorogov-Smirnov test. To account for the large degrees of freedom associated with comparing distributions and to prevent false positives, we decreased α to 0.001.

**Figure S1:**
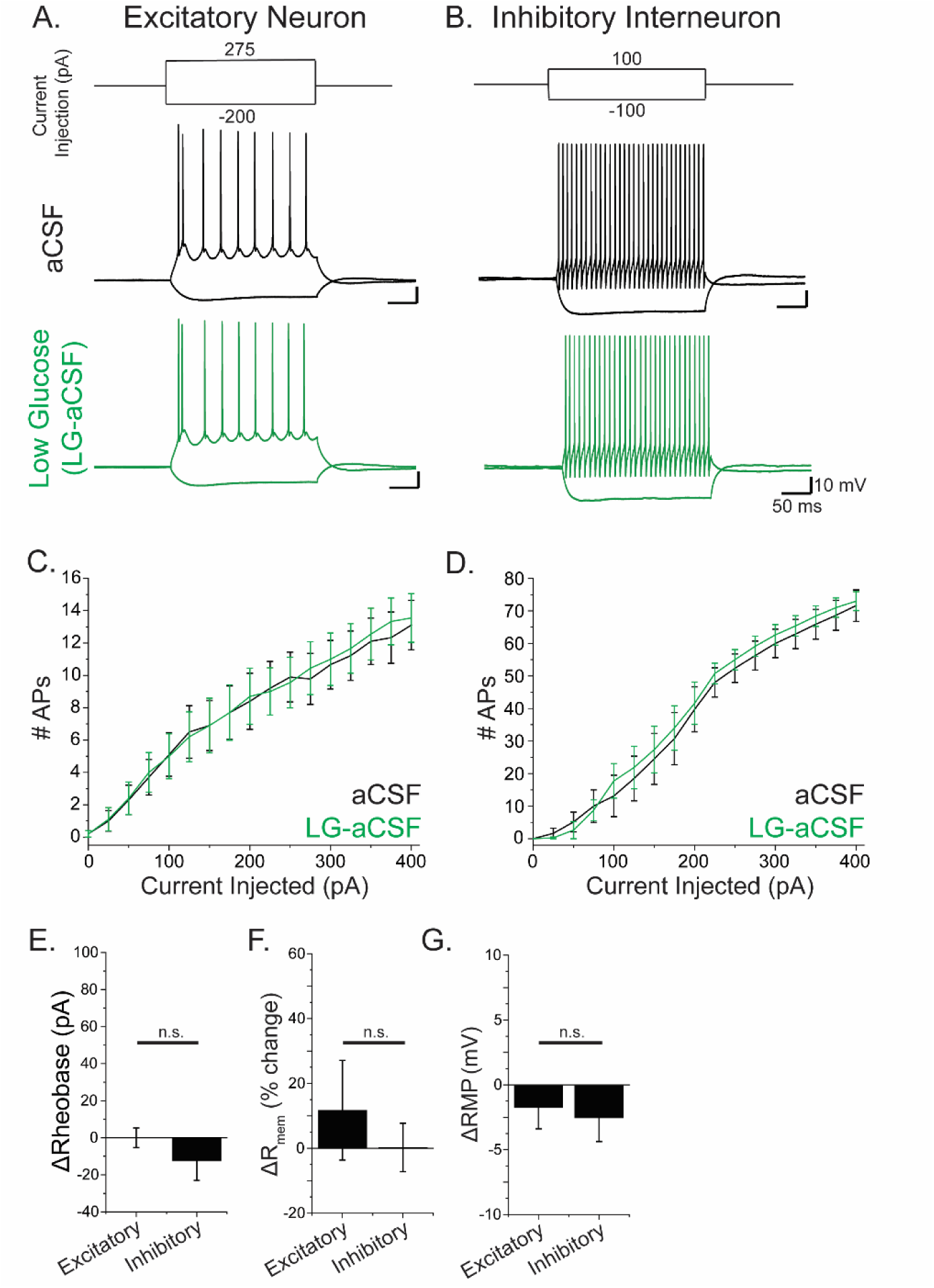
Treatment with Low Glucose (LG)-aCSF did not affect the excitability of excitatory or inhibitory neurons. **A-B.** Representative traces following current injection into Layer V cortical excitatory pyramidal neurons **(A)** or interneurons **(B)** before (black) or after (green) treating the cortical slice with LG-aCSF for 10 minutes. **C-D.** Input-output curves in excitatory (C) or inhibitory (D) neurons. **E.** ΔRheobase in excitatory and inhibitory neurons in LG-aCSF relative to baseline. **F.** % change of membrane resistance in excitatory and inhibitory neurons in LG-aCSF. **G.** Change in resting membrane potential (RMP) in excitatory and inhibitory neurons in LG-aCSF. (Error bar = SEM; n.s. not significant)

**Figure S2:**
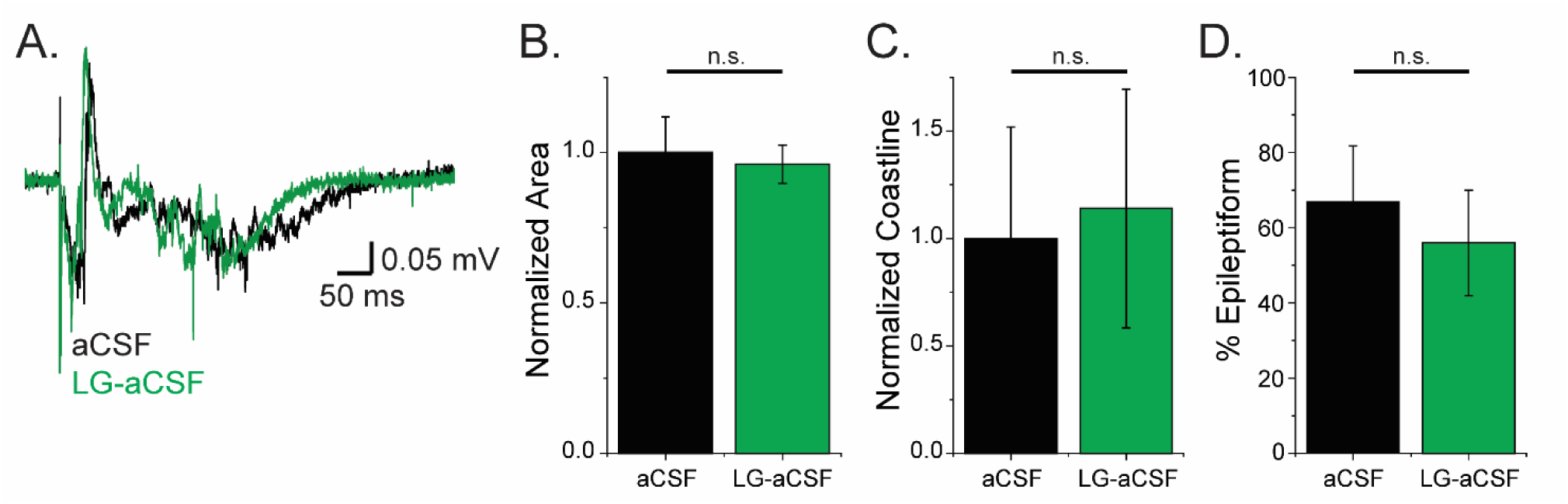
Low glucose (LG) conditions do not attenuate epileptiform activity following CCI. **A.** Representative stimulus-evoked field potentials in an acute cortical slice 3-5 weeks following CCI surgery. **B-C.** Area (B) and coastline (C) measurements from fEPSPs from CCI-injured animals with or without 30 minutes of LG-aCSF. **D.** The percentage of sweeps exhibiting epileptiform activity in slices from CCI animals with or without LG-aCSF treatment. (Error bar = SEM; n.s. not significant)

**Figure S3:**
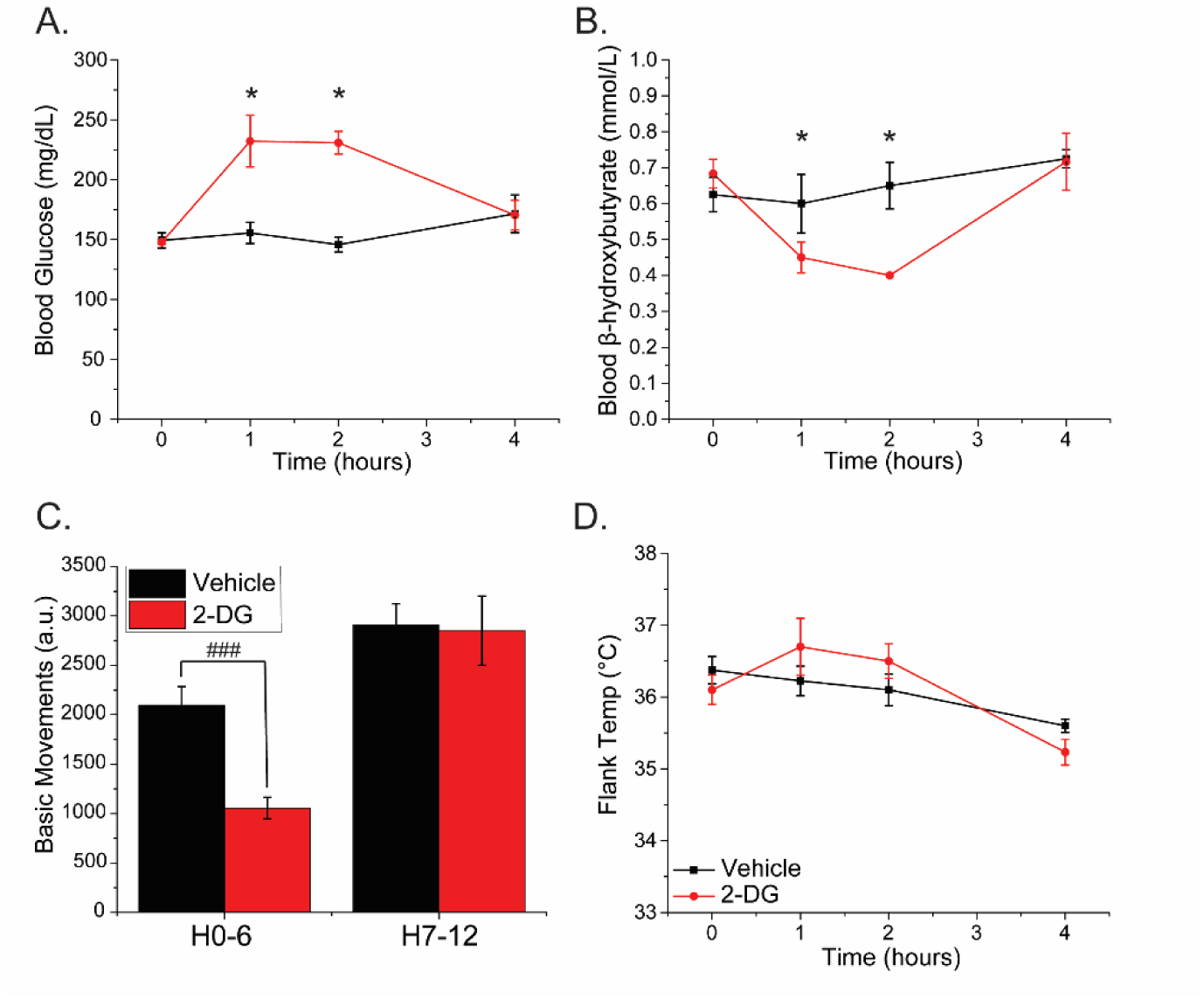
2-DG has transient effects on blood glucose and locomotor activity. **A.** Blood glucose in fasted mice after a single vehicle or 2-DG injection. **B.** Blood ketone body levels (as measured by β-hydroxybutyrate) after vehicle or 2-DG injection. **C.** Basic locomotor activity in vehicle- or 2-DG-treated animals. **D.** Flank temperature after a single vehicle or 2-DG injection. (Error bar = SEM; LMM: * t > ±1.96, effect: 2-DG at each time point; 2-sample t-test: ###p < 0.001)

**Figure S4:**
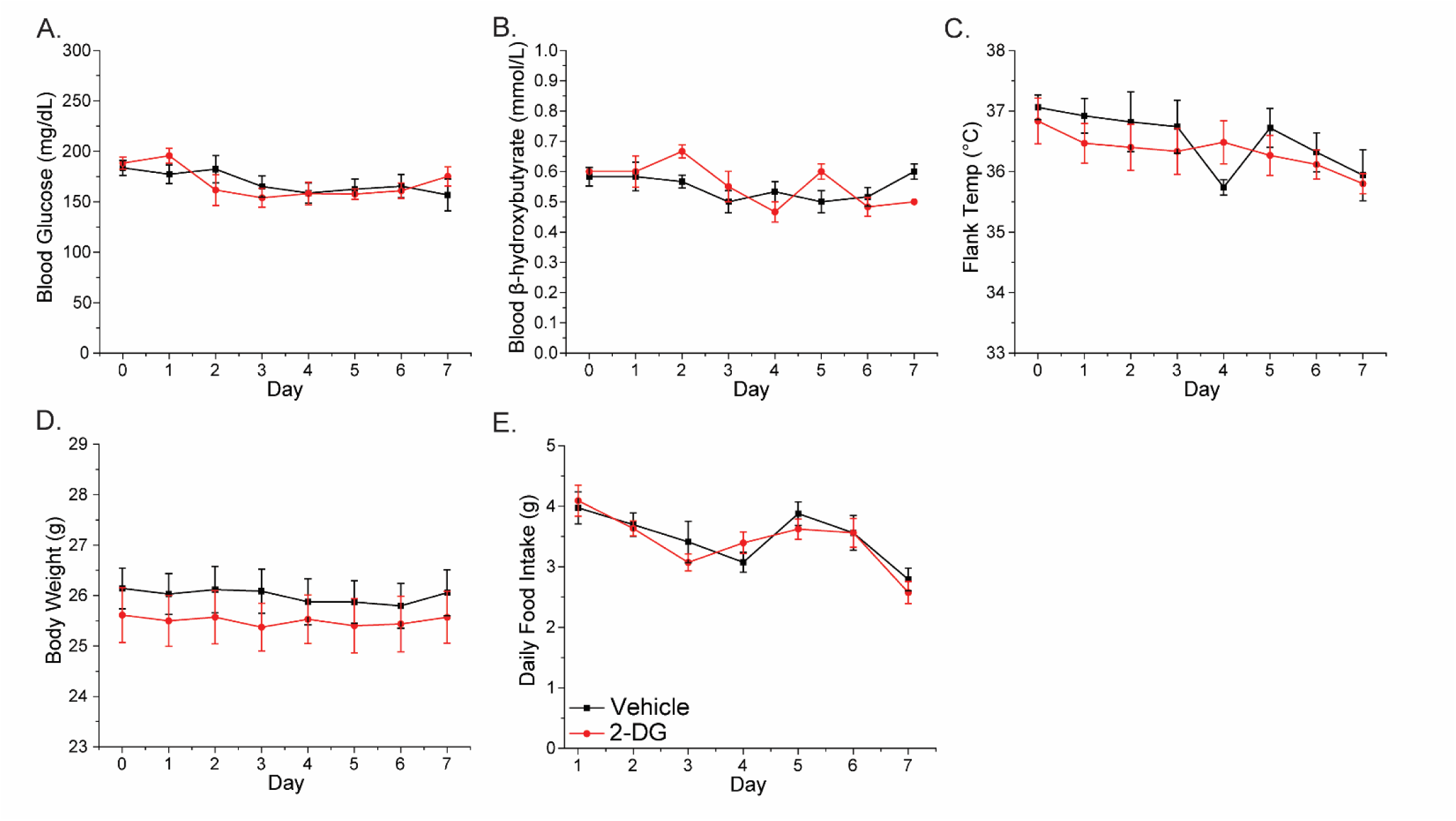
2-DG has no effect on daily blood glucose, blood β-hydroxybutyrate, temperature, body weight, or food intake during a week-long dosing regimen. Animals were injected daily with 2-DG or vehicle. Shown are blood glucose (**A**), β-hydroxybutyrate (**B**), flank temperature (**C**), body weight (**D**), and daily food intake (**E**) measured immediately prior to each daily injection. (Error bar = SEM)

**Table S1.**
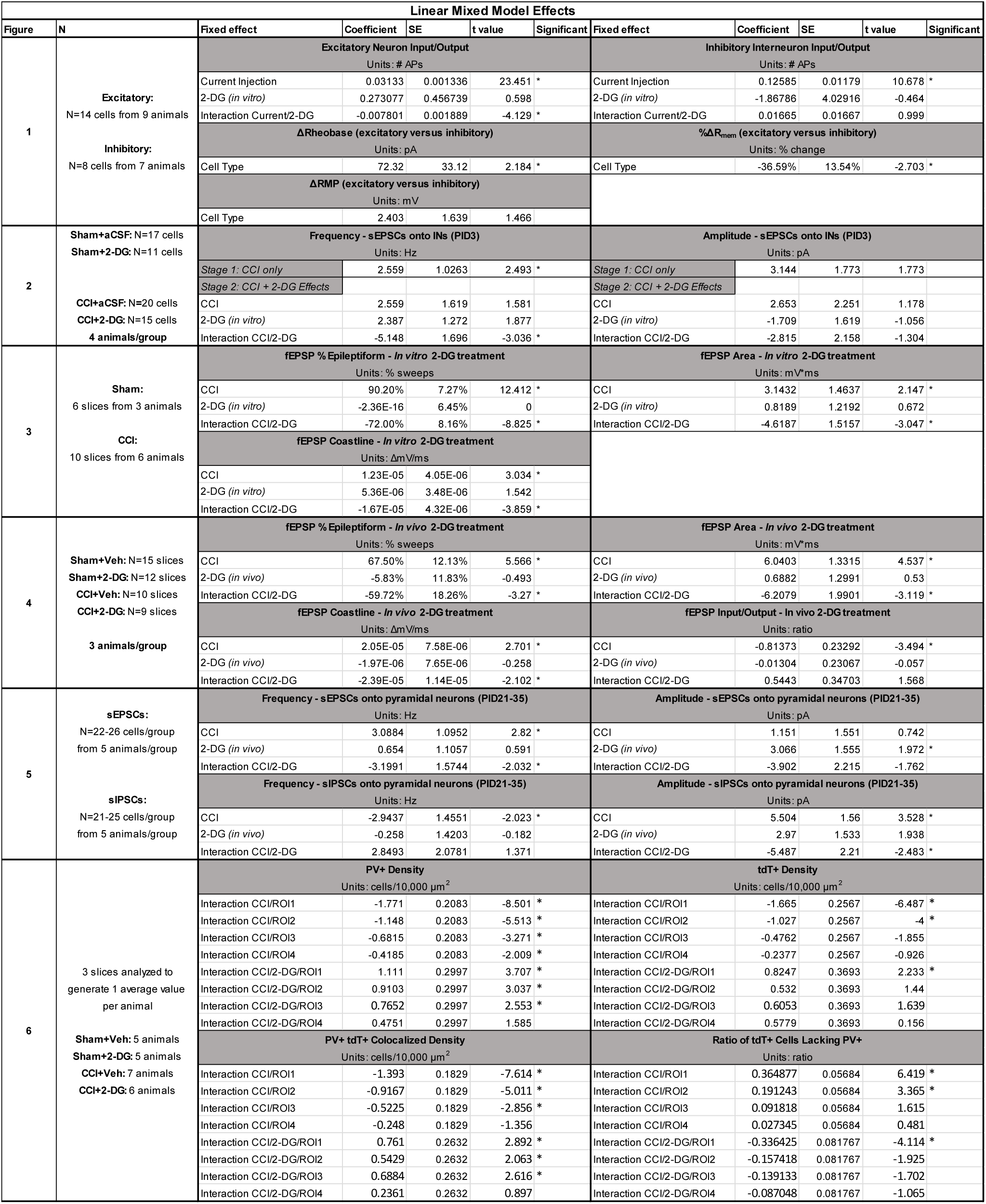
Linear mixed model results, organized by figure.

**Table S2.** Type III ANOVA results.

| Type III Analysis of Variance with Satterthwaite's Method |  |  |  |
| --- | --- | --- | --- |
| Data | Effect/Interaction | p value | Significant |
| PV+ Density | CCI | 1.83E-04 | * |
|  | 2-DG | 0.074809 |  |
|  | ROI | 1.27E-11 | * |
|  | CCI/ROI | 5.83E-15 | * |
|  | 2-DG/ROI | 1.21E-04 | * |
|  | CCI/2-DG | 0.058903 |  |
|  | CCI/2-DG/ROI | 0.002792 | * |
| tdT+ Density | CCI | 6.07E-09 | * |
|  | 2-DG | 0.35164 |  |
|  | ROI | 1.02E-06 | * |
|  | CCI/ROI | 1.48E-11 | * |
|  | 2-DG/ROI | 0.09617 |  |
|  | CCI/2-DG | 0.10985 |  |
|  | CCI/2-DG/ROI | 0.11161 |  |
| PV+ tdT+ Colocalized Density | CCI | 0.0037177 | * |
|  | 2-DG | 0.0729436 |  |
|  | ROI | 9.72E-09 | * |
|  | CCI/ROI | 7.91E-15 | * |
|  | 2-DG/ROI | 2.34E-04 | * |
|  | CCI/2-DG | 0.101366 |  |
|  | CCI/2-DG/ROI | 0.0202878 | * |
| Ratio of tdT+ Lacking PV+ | CCI | 0.307487 |  |
|  | 2-DG | 0.067198 |  |
|  | ROI | 1.58E-06 | * |
|  | CCI/ROI | 2.78E-06 | * |
|  | 2-DG/ROI | 1.34E-06 | * |
|  | CCI/2-DG | 0.127384 |  |
|  | CCI/2-DG/ROI | 0.001373 | * |

